# Ontogenic Shifts in Cellular Fate Are Linked to Proteotype Changes in Mouse Hematopoietic Progenitor Cells

**DOI:** 10.1101/2020.07.08.193276

**Authors:** Maria Jassinskaja, Kristýna Pimková, Emil Johansson, Ewa Sitnicka, Jenny Hansson

## Abstract

The process of hematopoiesis is subject to extensive ontogenic remodeling that is accompanied by alterations in cellular fate both during normal development and upon malignant transformation. Although the functional differences between fetal and adult hematopoiesis are well established, the responsible molecular mechanisms have long remained largely unexplored at the proteomic level. Here, we have applied state-of-the-art mass spectrometry to gain deep coverage of the proteome of 100,000 fetal and adult lympho-myeloid multipotent progenitors (LMPPs), common lymphoid progenitors (CLPs) and granulocyte-monocyte progenitors (GMPs). Our analysis resulted in the identification and quantification of 4189 proteins, with over 200 proteins per cell type displaying differential expression between the fetus and the adult. The proteomic data demonstrate that features traditionally attributed to adult hematopoiesis are conserved across lymphoid and myeloid lineages, while generic fetal features are considerably more prominent in LMPPs and CLPs than in GMPs. Furthermore, we reveal molecular and functional evidence for a diminished granulocyte differentiation capacity in fetal LMPPs and GMPs relative to their adult counterparts, and show indications of a differential requirement of myosin activity for granulopoiesis in fetal and adult LMPPs. We have additionally identified the transcription factor Irf8 as significantly lower expressed in fetal relative to adult GMPs, and shown that its expression pattern correlates with an altered capacity for monocytic differentiation in the fetal cells. Collectively, our work represents a significant advancement in the understanding of the molecular programs that govern ontogenic differences in early hematopoiesis and mature blood cell production.

**Key points:**

- In-depth proteomics links intrinsic molecular programs to functional output of fetal and adult lineage-biased hematopoietic progenitors
- Myelopoiesis-associated molecular programs and myeloid differentiation capacity are subject to considerable ontogenic remodeling

## Introduction

The lifelong support of the blood system relies on a complex differentiation program of hematopoietic stem cells (HSCs) and their progeny. The process of hematopoiesis occurs in waves with distinguishable functional characteristics from embryonic development through adulthood^1-3^. In mice, the largest burst of fetal hematopoiesis occurs in the fetal liver (FL) at embryonic day 14.5 (E14.5)^4-6^. After birth and throughout the lifetime of the animal, hematopoiesis is instead maintained in the bone marrow (BM)^2^.

The molecular programs that govern the functional differences between fetal and adult hematopoiesis have predominantly been investigated at the transcript level^7-9^. However, protein-level characterization is essential to account for post-transcriptional regulation, which plays a vital role in the function of hematopoietic stem and progenitor cells (HSPCs) both under normal and non-homeostatic conditions^1,10-13^. We have previously resolved the fetal- and adult-specific proteomic features that are shared among early hematopoietic cells (Lin^−^ Sca-1^+^ cKit^+^ [LSK] HSPCs)^1^, but the molecular programs that orchestrate important functions related to differentiation capacity and lineage-bias during development remain unresolved. Although a discrete hierarchical organization of hematopoiesis is subject to debate^14^, mature lymphoid and myeloid cell development is widely considered to occur through the differentiation of HSCs via several downstream hematopoietic progenitor cells (HPCs), including a progenitor with lympho-myeloid potential (lympho-myeloid multipotent progenitor [LMPP]), which gives rise to progenitors largely restricted towards the lymphoid (common lymphoid progenitor [CLP]) or myeloid (granulocyte-monocyte progenitor [GMP]) lineage^15^. Lineage-biased HPCs represent an important target for investigation, as such cells can act as potent leukemia-initiating cells^16^. Furthermore, recent findings indicate that the peculiar difference in incidence of leukemias of the myeloid and lymphoid lineage in children and adults^17^ at least in part originates from a differential susceptibility to leukemic transformation in lineage-biased HPCs at distinct stages of ontogeny^18^.

Here, we report the first-ever characterization of ontogenic changes that occur in the proteome of immunophenotypic LMPPs, CLPs and GMPs. We have quantified over 4000 proteins and uncovered striking differences in expression of hundreds of proteins between the fetal and the adult cells. Through extensive functional assays we link the proteomic signatures to cellular phenotypes, and uncover previously unknown ontogenic differences in the molecular makeup and functionality of lineage-biased HPCs.

## Methods

### Mice

Wild-type C57Bl/6N mice purchased from Taconic Biosciences or bred in-house were used for all experiments. Animals were housed in individually ventilated cages (IVC) and provided with sterile food and water *ad libitum*. All experiments involving animals were performed in accordance with ethical permits approved by the Swedish Board of Agriculture.

### Flow cytometry and FACS

E14.5 FL and adult BM (ABM) cells were depleted for Ter119^+^ cells and lineage-positive cells, respectively, by MACS depletion (Miltenyi). Cells were surface-stained with fluorophore-conjugated antibodies and incubated with 7AAD briefly before analysis to exclude dead cells. All flow cytometry and flouresence-activated cell sorting (FACS) experiments were performed on BD instruments FACSAriaIIu, FACSAriaIII, LSRFortessa or LSRFortessa X-20. Data analysis was performed in FlowJo (BD).

### Proteome analysis

Cell pellets corresponding to 100,000 FACS-sorted cells were processed using in-StageTip (iST) NHS kit (PreOmics) in accordance with manufacturer’s protocol. Digested peptides were labeled using TMT6plex reagents (Thermo Scientific) and fractionated by a high-pH-reverse phase (HpH-RP) approach as previously described^19^ with some modifications. MS analyses were carried out on an Orbitrap Fusion Tribrid MS instrument (Thermo Scientific) equipped with a Proxeon Easy-nLC 1000 (Thermo Fisher) using a 120 min linear gradient separation and a data-dependent synchronous precursor scanning MS3 (SPS-MS3) acquisition method. MS raw data files were processed with Proteome Discoverer (version 2.2.0; Thermo Scientific). The derived peak list was searched using the Sequest HT node against the Swissprot mouse database (version 2017.07.05) together with commonly observed contaminants and reversed sequences for all entries. Proteins were quantified based on the average corrected TMT reporter ion intensities from two technical replicates per sample.

### *In vitro* differentiation assays

HPCs were seeded onto pre-established layers of OP9/OP9DL1 stroma cells. Cells were cultured in complete medium supplemented with cytokines promoting B/myeloid or T cell differentiation. For ROCK inhibition, cells were cultured together with H1152 (R&D systems) or sterile water. For suspension cultures, cells were cultured in complete medium supplemented with cytokines promoting monopoiesis. Megakaryocyte potential was assessed using the MegaCult culture system (StemCell Technologies) in accordance with manufacturer’s protocol.

### Statistical analysis

Statistical analysis of proteome data was performed using the Limma package in R/Bioconductor^20^ with multiple testing correction using Benjamini-Hochberg’s method. Proteins with an adjusted p value < 0.05 were considered to be differentially expressed. For all other experiments, differences between two groups were assessed by two-tailed Students’ t-test using Prism software version 8 (GraphPad). Error bars represent SD. ****p<0.0001, ***p<0.001, **p<0.01, *p<0.05 and ns=non-significant.

## Data sharing

The proteomics data have been deposited to the MassIVE repository.

## Results

### Immunophenotypic LMPPs, CLPs and GMPs show ontogeny-specific lineage output and differentiation kinetics

The immunophenotype and lineage-potential of murine adult LMPPs, CLPs and GMPs have been extensively studied^21-25^, whereas less is known about the cells’ counterparts during fetal life. Using the immunophenotypic definitions previously described for the adult hematopoietic hierarchy^21-24^, we first determined the occurrence of LMPPs (LSK Flt3^high^ CD150^−^), CLPs (Lin^−^ Sca-1^low^ cKit^low^ IL-7Ra^+^ Flt3^high^) and GMPs (Lin^−^ Sca-1^−^ cKit^+^ [LS^−^K] CD41^low^ CD150^−^ CD16/32^+^) in E14.5 FL and ABM (Figure 1A-C). The frequency of GMPs was 2-6% in both tissues, while LMPPs and CLPs were at an approximately ten-fold lower frequency for both tissues, comprising only 0.2-0.6% of the FL and ABM (Figure 1B).

**Figure 1.**
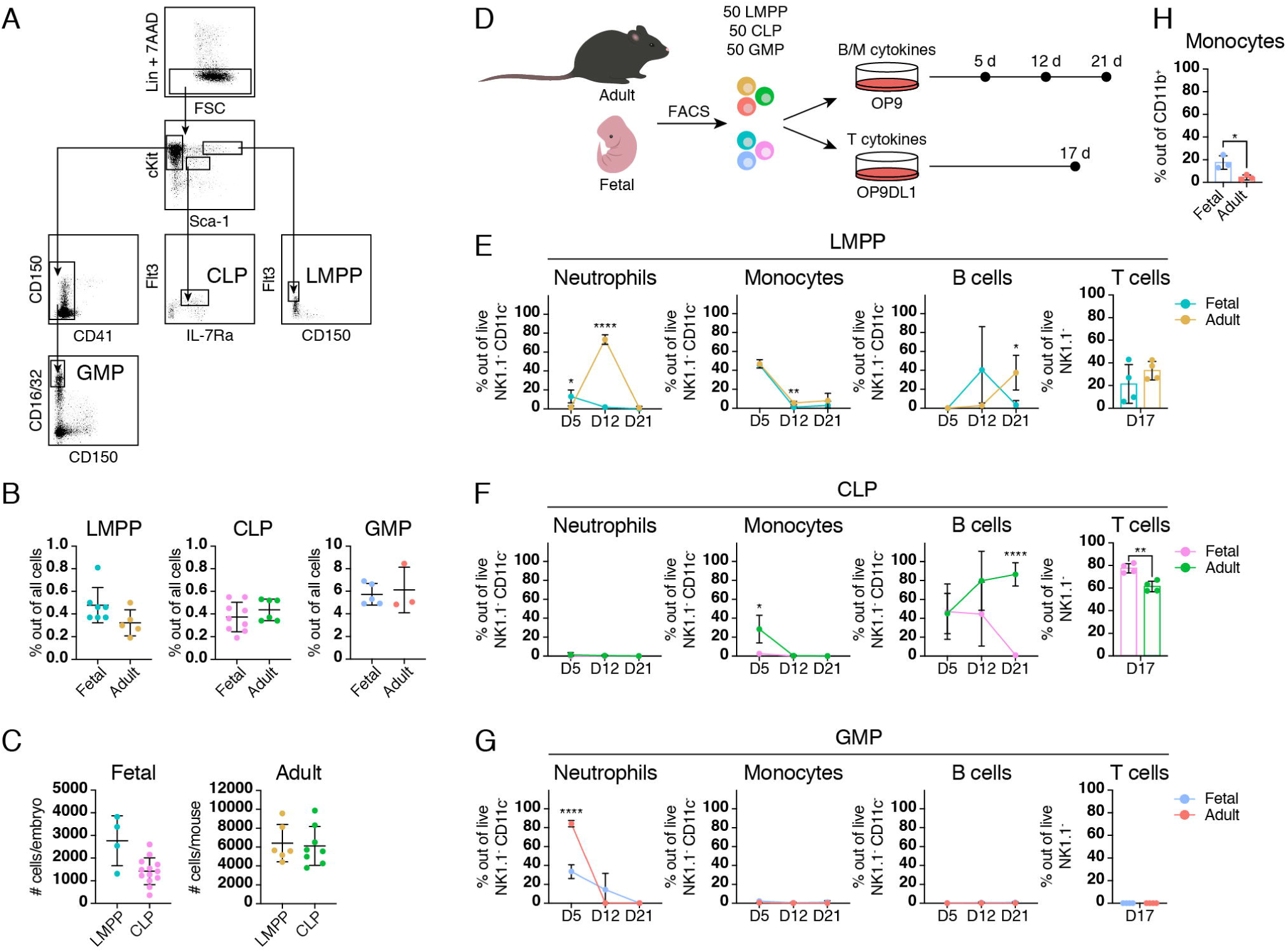
Fetal and adult HPCs show distinct lineage potential and differentiation kinetics. (A) Representative FACS plots showing the gating strategy for LMPPs (LSK Flt3^high^ CD150^−^), CLPs (LS^low^K^low^ Flt3^+^ IL7R^+^) and GMPs (LS^−^K CD41^low^ CD150^−^ CD16/32^+^). (B) Frequency of LMPPs, CLPs and GMPs in lineage-depleted E14.5 FL and ABM. (C) Number of LMPPs and CLPs present in one embryo and one adult mouse (hind limbs, fore limbs, spine and sternum) extrapolated from FACS sorting. (D) *In vitro* differentiation workflow for evaluation of fetal and adult LMPP, CLP and GMP lineage-potential. (E-G) Frequency of neutrophils (NK1.1^−^ CD11c^−^ CD11b^+^ Ly6G^+^), monocytes (NK1.1^−^ CD11c^−^ CD11b^+^ CD115^+^), B cells (NK1.1^−^ CD11c^−^ CD11b^−^ CD19^+^ B220^+^) and T cells (NK1.1^−^ Thy1^+^ CD25^+^ and/or NK1.1^−^ CD4^+^ CD8^+^) at different time points in wells seeded with fetal and adult LMPPs (E), CLPs (F) and GMPs. Data are depicted as mean of three biological replicates. (G). Data shown are from 4 biological replicates. (H) Frequency of monocytes after 7 days suspension culture of fetal and adult GMPs. Error bars represent mean ± SD. *p<0.05, **p>0.01, ***p<0.001 and ****p<0.0001.

We next evaluated *in vitro* lineage potential of the six cell populations under conditions promoting either B and myeloid or T cell fate (Figure 1D-H, Supplemental Figure 1A). Both fetal and adult LMPPs showed myeloid as well as B and T cell potential (Figure 1E), as previously described^23^. Compared to fetal LMPPs, adult LMPPs produced very high frequencies of neutrophils after 12 days of culture (Figure 1E), indicating a greatly elevated capacity to generate granulocytes. From CLPs, B cell output was observed already after 5 days from both fetal and adult cells, and T cells were potently produced in parallel cultures (Figure 1F). We additionally detected monocytes in adult CLP wells. Critically, fetal CLPs showed exceedingly low monocyte production (Figure 1F), indicating a stronger lineage-restriction. GMPs from fetus and adult exclusively generated myeloid cells (Figure 1G). Surprisingly, we observed almost no monocytes in wells seeded with GMPs, despite our system supporting monocytic differentiation from other progenitor subtypes (Figures 1E and 1F). Suspension culture of GMPs in the presence of macrophage-colony stimulating factor (M-CSF) demonstrated that both fetal and adult GMPs have the potential to generate monocytes, and that this potential is significantly higher in fetal relative to adult cells (Figure 1H). Collectively, our data show ontogeny-specific variation in the differentiation kinetics and intrinsic lineage-bias of fetal and adult HPCs in the form of a stronger myeloid potential in adult LMPPs and CLPs relative to their fetal counterparts, as well as a differential output of neutrophils and monocytes from fetal and adult GMPs.

### The proteomic landscapes of LMPPs, CLPs and GMPs undergo extensive ontogenic remodeling

Having confirmed differential lineage-potential of fetal and adult HPCs, we moved on to determine the proteomic profiles of these cells. We subjected 100,000 FACS-purified LMPPs, CLPs and GMPs (Figure 1A and Supplemental Figure 1A) from E14.5 FL and ABM in three biological replicates to our proteomic workflow (Figure 2A, Supplemental Figure 1B and 1C). We identified 4,189 proteins, out of which 4,021 proteins were quantified in all six cell populations (Figure 2B, Supplemental Figure 1D and 1E and Supplemental Table 1).

**Figure 2.**
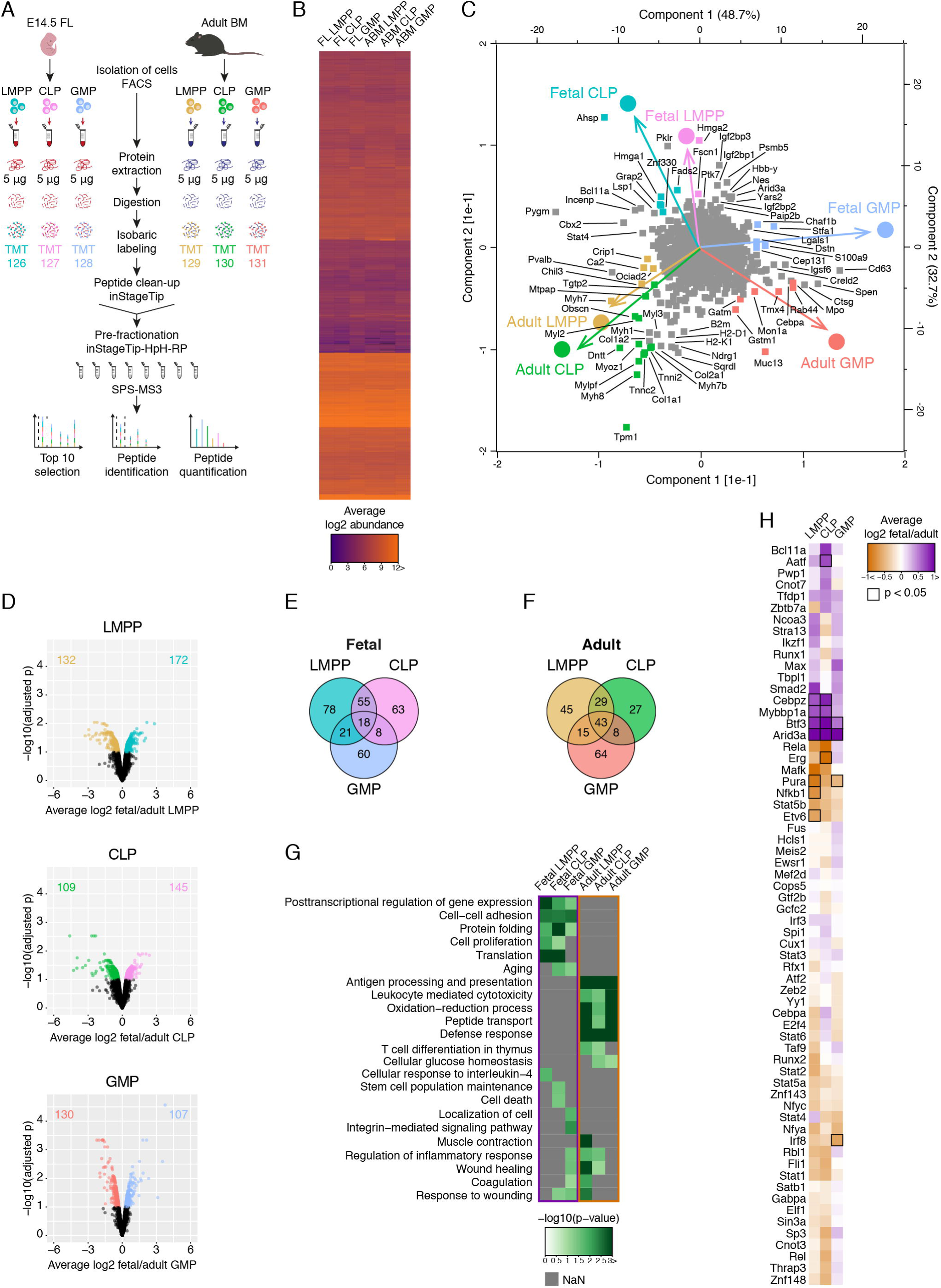
Ontogenic remodeling of the proteome of LMPPs, CLPs and GMPs. (A) Workflow for proteomic analysis of fetal and adult LMPPs, CLPs and GMPs. (B) Heatmap depicting the average abundance of 4032 proteins quantified in at least one cell population. iST-based methods were utilized for sample preparation^56^ and HpH-RP pre-fractionation^19^. Samples were labeled with isobaric tags and analyzed using an SPS-MS3 method^57^. (C) PCA of relative protein expression. Colored proteins are likely contributing most to the separation. (D) Statistical analysis of proteins differentially expressed between fetal and adult LMPPs, CLPs and GMPs. Proteins with a significantly higher expression (adjusted p-value < 0.05) are shown in color. (E, F) Overlap of proteins significantly higher expressed in fetal (E) and adult (F) cells. (G) Heatmap showing selected results from GO analysis of biological processes enriched in differentially expressed proteins. Only processes with a p-value < 0.05 are shown. Processes that were not detected are depicted in grey. See Supplemental Figure 1H for a full depiction of enriched biological processes. (H) Heatmap depicting the relative expression of proteins classified as transcription factors by PANTHER^58^ and validated by TRRUST^59^. Proteins differentially expressed between fetus and adult are indicated by a black frame. All expression data represent the average of three biological replicates.

Principal component analysis (PCA; Figure 2C) segregated CLPs as well as LMPPs on ontogenic stage, while these two lymphoid-competent progenitors positioned close to each other for FL and ABM, respectively, indicating a particularly strong expression of fetal- and adult-specific features in LMPPs and CLPs. Conversely, the rather close positioning of fetal and adult GMPs points towards stronger ontogenic conservation in these cells. As expected, proteins contributing to generic fetal signatures included known fetal-enriched proteins such as transcription factor (TF) Arid3a^26^ and insulin-like growth factor-binding proteins (Igf2bp1, Igf2bp2 and Igf2bp3)^1,27^. Adult characteristics covered proteins related to antigen presentation (e.g. H2-K1, H2-D1 and B2m) which we have previously shown to be higher expressed in adult compared to fetal HSPCs^1^ (Supplemental Figure 1F). The fetal LMPP signature appeared strongly influenced by Lin28b-target Hmga2^28^, while lymphopoiesis-associated proteins such as Grap2^29^, Bcl11a^30^ and Lsp1^31^ contributed to the signature of fetal CLPs. Intriguingly, another protein causative to the separation of fetal CLPs from the other cell populations was alpha-hemoglobin stabilizing protein (Ahsp). This molecular chaperone has a well-documented role in erythropoiesis^32^, but its function within the context of lymphopoiesis has not been described. Among proteins contributing to the fetal GMP signature, we identified the pro-inflammatory protein S100a9, and, interestingly, megakaryocyte protein Stfa1^33^. The adult LMPP signature included Stat3-activator Ociad2^34^, while adult lymphopoiesis-specific enzyme Dntt^1^ contributed strongly to the signature of adult CLPs. Surprisingly, the proteomic signatures of adult LMPPs and CLPs were strongly influenced by several members of the myosin, tropomyosin and troponin family of proteins (e.g. Mylpf, Myoz1, Tpm1, Myh7, Myh7b and Myl2), a to our knowledge previously unknown hallmark of adult versus fetal lymphoid-competent progenitors. Myelopoiesis-associated proteins (e.g. Cebpa, Mpo, Rab44)^35,36^, as well as redox proteins Tmx4 and Gstm1 could be found to represent features of adult GMPs.

### Differential protein expression between fetal and adult HPCs is indicative of ontogenic changes in functionality

In contrast to up-stream HPCs, which display a considerably higher proteome complexity in adult relative to the fetus^1^, HPCs showed rather equal distribution of differentially expressed proteins (Supplementary Figure 1G), indicating a similar level of proteome complexity in these cells across ontogeny. Statistical analysis of proteins quantified in at least two replicates identified 304, 254 and 237 proteins as differentially expressed between fetal and adult LMPPs, CLPs and GMPs, respectively (adjusted p-value < 0.05; Figure 2D). As already indicated by PCA (Figure 2C), the number of differentially expressed proteins point to GMPs as the least ontogenically distinct cell type. Proteins significantly higher expressed in fetal LMPPs and CLPs compared to their adult counterparts showed a high overlap (55 proteins), whereas only 18 proteins were shared among all three fetal cell types (Figure 2E). The corresponding numbers in the adult were 29 and 43 proteins, respectively (Figure 2F), indicating that adult-specific features are conserved across different hematopoietic lineages, whereas fetal-specific features diverge between lymphoid-competent and myeloid-restricted progenitors.

Gene ontology (GO) enrichment analysis showed that common features of proteins higher expressed in fetal HPCs are related to translation (mainly ribosomal proteins) and cell-cell adhesion (e.g. Lgals9, Lgals1, S100a8 and S100a9) (Figure 2G and Supplemental Figure 1H). Proteins higher expressed in adult HPCs were instead enriched for immune response-related processes and oxidation-reduction processes (mainly glycolytic proteins and proteins involved in glutathionylation), similar to adult HSPCs^1,7^. In line with the adult LMPP-signature in our PCA (Figure 2C), “Muscle contraction” was enriched in proteins significantly higher expressed in adult LMPPs and included Myh1, Myl1 and Myh4, among others. Interestingly, fetal GMPs were the only cell type in the fetus where differentially expressed proteins showed an enrichment for multiple processes related to inflammation (e.g. “Regulation of inflammatory response”, “Coagulation” and “Wound healing”; Figure 2G and Supplemental Figure 1H), including proteins such as Apoe, Plek, Vcl and Ptk7. This suggests that fetal GMPs are more strongly intrinsically poised to react to inflammatory insult compared to their adult counterpart.

Analysis of TFs revealed differential expression of known regulators of hematopoiesis (Arid3a, Erg, Nfkb1, Etv6 and Irf8), as well as several TFs with poorly described or undefined functions in early blood cell development (Aatf, Cebpz, Mybbp1a, Btf3 and Pura; Figure 2H and Supplemental Figure 2A). In addition to confirming the requirement of Erg in adult B lymphopoiesis^37-39^, the expression profile of this TF excludes it as a major driver of fetal lymphopoiesis (Supplemental Figure 2A). Intriguingly, Btf3 – a TF with no defined role in hematopoiesis – followed the same expression pattern as the known fetal-enriched TF Arid3a^26^ (Figure 2H), indicating its potential role as a regulator of fetal-specific features of hematopoiesis. Btf3 has been suggested to act as an inhibitor of the Nf*κ*b signaling pathway^40^, of which we found a major part to be downregulated in fetal compared to adult HPCs (Supplemental Figure 2B). Importantly, Mybbp1a, which we identified as significantly higher expressed in fetal LMPPs and CLPs compared to their adult counterparts (Figure 2H), has also been implicated as a Nfκb-repressor^41^, suggesting that several mechanisms function to dampen inflammatory responses driven by Nfκb in fetal HPCs.

### Inhibition of myosin activity enhances granulopoiesis from adult LMPPs

One of the main proteomic features distinguishing fetal and adult LMPPs was a significantly higher expression of many members of the myosin, tropomyosin and troponin family of proteins in the adult cells (Figure 3A). Considering the strong ontogenic expression differences and the previously reported roles of this protein family in acute myeloid leukemia (AML)^42^ as well as inflammation^43^ and emergency hematopoiesis^11^ (Figure 3B), we hypothesized that fetal and adult LMPPs may exhibit a differential dependency on myosin activity for expansion and/or myeloid differentiation. We targeted the actin/myosin motor by interfering with myosin light chain (MLC) phosphorylation via inhibition of Rho kinase (ROCK)^42,44^. In two independent experiments, reduction of myosin activity modestly enhanced granulopoiesis in adult, but not fetal, LMPPs (p=0.15 and p=0.03; Figure 3C), supporting an adult-specific role in hematopoiesis. We also found a reduced expansion of adult LMPPs in one of the experiments (Figure 3D). We next investigated the myeloid output of fetal and adult LMPPs upon induction of inflammatory stress with interferon alpha (IFNα. We did not observe any difference in granulocyte output between fetal and adult LMPPs (Supplemental Figure 2C), indicating that emergency granulopoiesis is regulated independently of homeostatic myosin expression levels in LMPPs. Collectively, our results suggest an ontogeny-specific role of myosins in regulating granulocytic output from LMPPs (Figure 3E).

**Figure 3.**
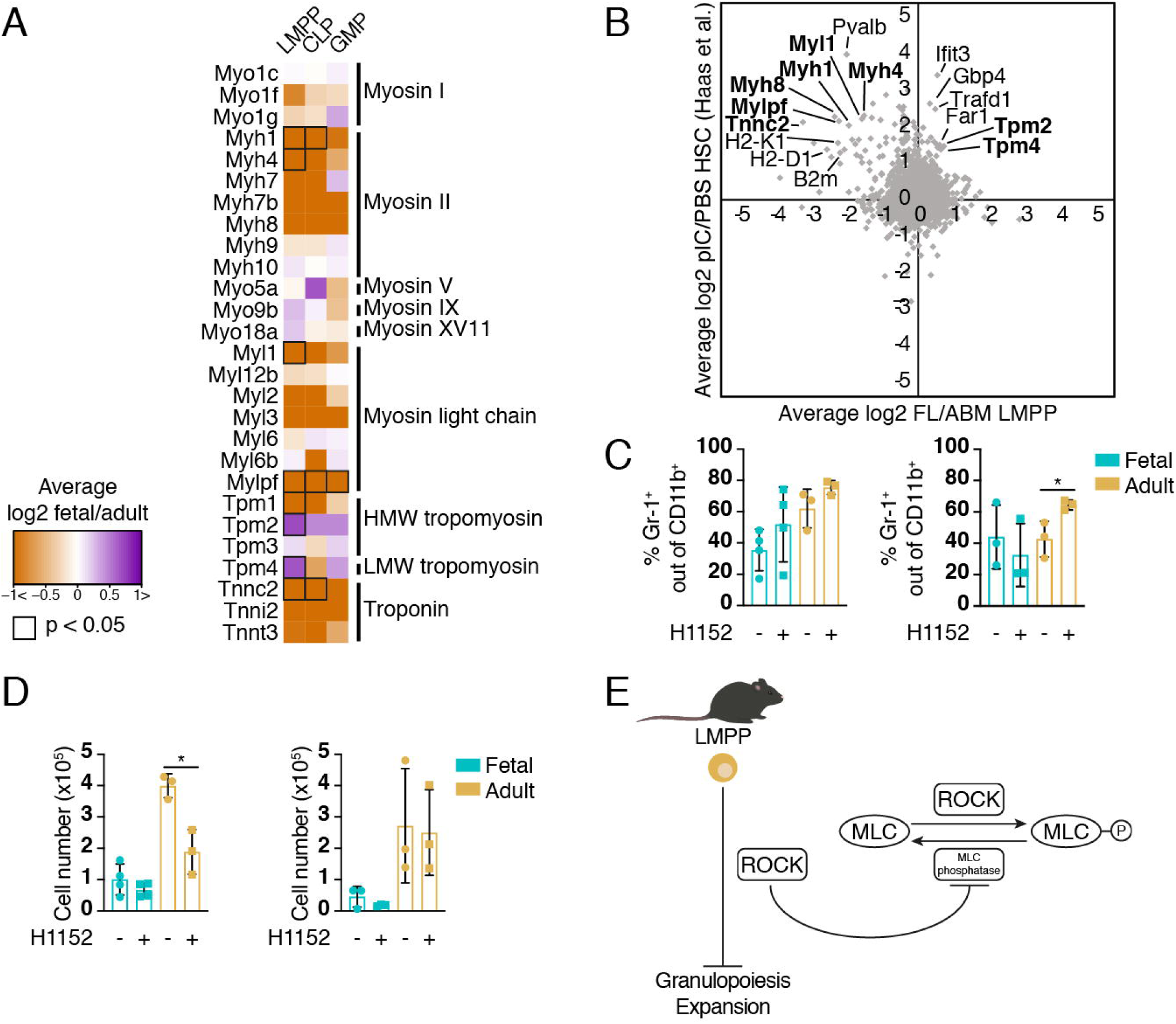
Fetal and adult LMPPs exhibit differential sensitivity to ROCK inhibition. (A) Average expression of proteins belonging to the myosin, myosin light chain, tropomyosin and troponin family of proteins in fetal and adult LMPPs, CLPs, and GMPs. Differentially expressed proteins are indicated by a black frame. HMW = high molecular weight, LMW = low molecular weight. (B) Correlation between fetal/adult LMPP protein ratios and poly(I:C)/PBS-treated HSC protein ratios^11^. Proteins belonging to the myosin, tropomyosin or troponin family of proteins are indicated in bold. (C, D) Frequency of Gr-1^+^ granulocytes (C) and cell count (D) following treatment with H1152. The two panels represent two different experiments. Error bars are ± SD. *p<0.05.

### Proteotype of fetal and adult HPCs accurately predicts ontogenic changes in myeloid potential while MegE potential is uncoupled from protein signature

We next set out to investigate the relationship between differentially expressed proteins and lineage-bias. We mapped differentially expressed proteins to transcriptional profiles of various subsets of mouse adult hematopoietic cells^45^ (Figure 4A-4C and Supplemental Figure 3A), as well as investigated expression of individual proteins known to be involved in lineage-commitment (Figure 4D). Using this approach for proteins differentially expressed between CLPs and GMPs in the fetus and adult showed, as expected, a high similarity between the profiles with a clear association of proteins higher expressed in CLPs and GMPs to lymphoid and myeloid cell subsets, respectively (Supplemental Figure 3A). In addition, a high correlation was observed between fetal and adult CLP versus GMP protein expression signatures (Supplemental Figure 3B and 3C), suggesting that the main features that discriminate lymphoid from myeloid cell fate are conserved during ontogeny. Notably, despite sustained lineage commitment signatures, we found strong ontogeny-specific differences in the expression of myelopoiesis-associated proteins, where fetal GMPs showed significantly lower expression of several proteins linked to myeloid commitment, such as Irf8, Mpo, and Elane (Figure 4D), compared to adult GMPs. In line with our *in vitro* differentiation data (Figure 1E and 1F), proteins higher expressed in adult LMPPs and CLPs associated more strongly with mature myeloid cell subsets compared to the corresponding fetal signatures (Figure 4A and 4B). Among individual proteins known to be involved in myelopoiesis, Cst7 was found significantly higher expressed in adult CLPs compared to fetal CLPs (Figure 4D). Intriguingly, compared to the adult, fetal LMPP and CLP signatures showed a high association with the erythroid lineage (Figure 4A and 4B). We found no capacity for erythroid differentiation in either of these cell types (Supplemental Figure 3D), suggesting that fetal LMPPs and CLPs share features with adult erythroid progenitors that are not directly linked to red blood cell production. Indeed, among the proteins significantly upregulated in fetal LMPPs and CLPs, translation- and/or proliferation-associated proteins Rrm1, Heatr3, Eif2s3x, Larp4 and Birc5 have high transcript expression in adult CFU-Es^45^.

**Figure 4.**
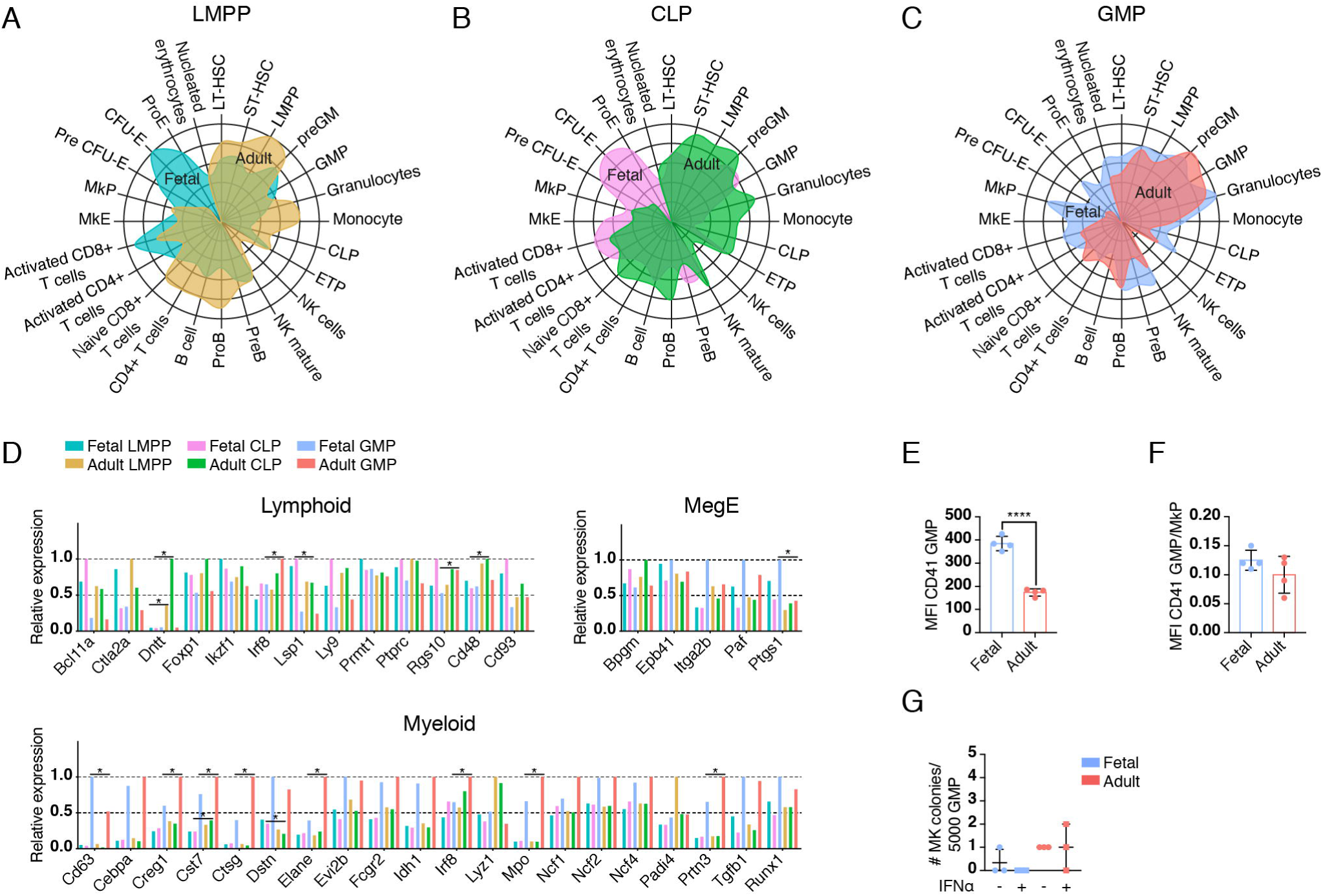
Proteomic profiles predict differential functionality and lineage potential of fetal and adult HPCs. (A-C) Radar plots depicting the association of differentially expressed proteins in LMPPs (A), CLPs (B) and GMPs (C) with known transcriptional profiles of murine hematopoietic cell subsets. (D) Relative expression of proteins associated with the lymphoid myeloid and megE lineage. Data represent the average of three biological replicates. *p<0.05. (E) CD41 expression on fetal and adult GMPs. (F) CD41 expression on fetal and adult GMPs relative to MkPs. (G) MK potential in fetal and adult GMPs with or without IFNα treatment. MFI = median fluorescent intensity. Error bars are ± SD. *p<0.05 and ****p<0.0001.

In accordance with proteins significantly upregulated in fetal GMPs showing enrichment for biological processes related to inflammation and coagulation (Figure 2G and Supplemental Figure 1H), the proteomic profile of these cells also showed a pronounced association with megakaryocyte progenitors (MkPs) compared to that of their adult counterpart (Figure 4C). Strikingly, expression of MegE-associated proteins Itga2b (CD41)^24^, Paf^46^ and Ptgs1^11^ was elevated in fetal relative to adult GMPs (Figure 4D), in line with our other observations of a potential retainment of MegE potential in fetal myeloid progenitors (Figure 2G, Figure 4C and Supplemental Figure 1H). Analysis of cell surface CD41 levels confirmed a significantly higher expression in fetal compared to adult GMPs (Figure 4E). Importantly, CD41 expression on fetal GMPs was still approximately 85% lower than that observed on MkPs (Figure 4F). To exclude the possibility that CD41^+^ fetal cells represent primitive erythro-myeloid progenitors (EMPs) persisting from earlier waves of embryonic hematopoiesis^47^, we assessed the macrophage potential of the fetal cells. Although the frequency of granulocytes produced from CD41^−^ relative to CD41^+^ fetal GMPs was elevated, both fractions generated equally low numbers F4/80^+^ macrophages (Supplemental Figure 3E), confirming that CD41^+^ fetal GMPs represent a population separate from EMPs. Because the frequency of immunophenotypic LS^−^K CD150^+^ CD41^+^ MkPs is greatly reduced in the FL relative to the ABM (Supplemental Figure 3F), we next investigated whether GMPs may act as an alternative source of megakaryocytes during fetal life. Surprisingly, we found very limited megakaryocyte production from fetal as well as adult GMPs under both homeostatic and stressed conditions (Figure 4G). In summary, while the proteomic signatures of HPCs correlate with ontogenic alterations in myeloid potential, the shared proteomic features of fetal GMPs with MkPs, including an elevated expression of CD41, do not have a strong impact on the cells’ ability to produce megakaryocytes.

### Ontogeny-specific differences in Irf8 expression correlate with an altered capacity for monocyte differentiation in GMPs

An intriguing finding from our analysis of TFs and myelopoiesis-associated proteins was a significantly elevated expression of Irf8 in adult relative to fetal GMPs (Figure 2H and Figure 4D). Loss of Irf8 has a severe impact on the generation of mature monocytes, and is sufficient to produce a chronic myelogenous leukemia-like disease in mice^48-50^. Considering this remarkably strong phenotype, we decided to explore the capacity for monocytic differentiation in fetal and adult GMPs. The frequency of produced myeloid cells was low from both fetal and adult GMPs, but significantly higher from adult compared to fetal cells (Supplemental Figure 3G). In contrast to what would be expected considering that Irf8 promotes monocytic cell fate, fetal GMPs produced significantly higher frequencies of monocytes than did adult GMPs, whereas the opposite was true for neutrophil generation (Figure 5A). However, upon analysis of expression of Ly6C, a marker for inflammatory monocytes^51^, we found that the ratio of Ly6C^+^ to Ly6C^−^ monocytes was approximately 50% lower in fetal-relative to adult-derived cells (Figure 5B). Additionally, we observed an accumulation of CD11b^−^ Ly6C^+^ CD115^+^ cells immunophenotypically analogous to monocyte progenitors (MPs)^49^ in fetal GMP wells (Supplemental Figure 3H). Furthermore, Ly6C^+^ monocytes derived from fetal GMPs showed a significantly lower Ly6C surface expression relative to those produced from adult GMPs (Figure 5C), suggesting incomplete monocytic maturation and a partial block in monocyte differentiation in fetal GMPs. Importantly, this phenotype shares many similarities with that previously observed in *Irf8*^*−/−*^ mice^49,50^.

**Figure 5.**
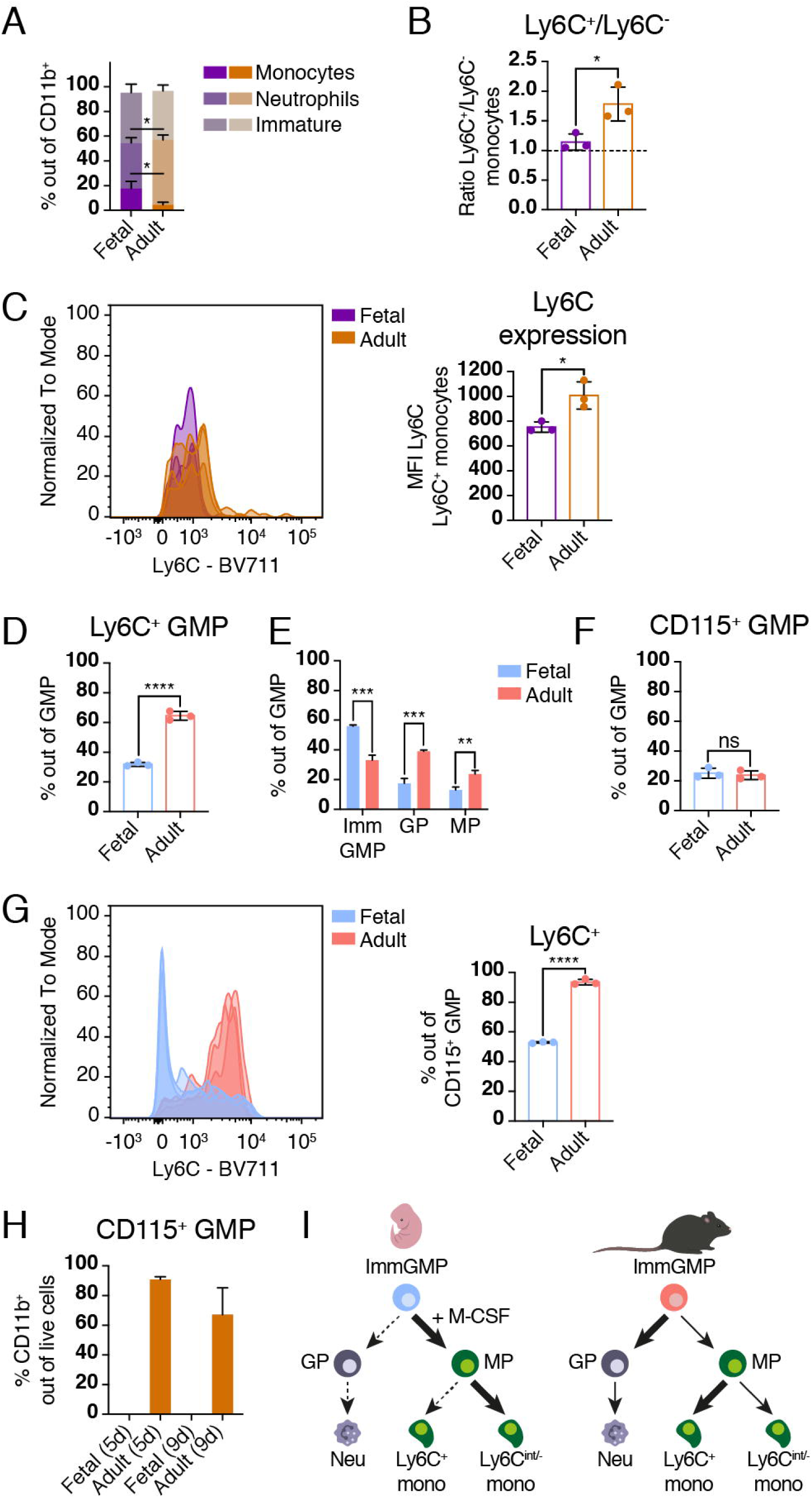
Differential expression of Irf8 correlates with ontogenic alterations in monocyte differentiation. (A) Frequency of neutrophils, monocytes and immature myeloid cells (NK1.1^−^ CD11c^−^ CD11b^+^ Ly6G^−^ CD115^−^) after 7 days suspension culture of fetal and adult GMPs. Data are depicted as mean of three biological replicates. (B, C) Ratio of Ly6C^+^ to Ly6C^−^ monocytes (B) and Ly6C expression on Ly6C^+^ monocytes (C) derived from *in vitro* cultured fetal and adult GMPs. (D) Frequency of Ly6C^+^ cells within the fetal and adult GMP pool. (F) Frequency of Ly6C^−^ CD115^−^ immature GMPs, Ly6C^+^ CD115^−^ GPs and Ly6C^+^ CD115^+^ MPs within the fetal and adult GMP pool. Data are depicted as mean of three biological replicates. Imm = immature. (F) Frequency of CD115^+^ cells within the fetal and adult GMP population. (G) Ly6C expression on fetal and adult CD115^+^ GMPs. (H) Frequency of CD11b^+^ cells derived from fetal and adult CD115^+^ GMPs. MFI = median fluorescent intensity. Error bars are ± SD. *p<0.05, **p>0.01, ***p<0.001, ****p<0.0001 and ns=non-significant.

Because it has been suggested that Irf8 has its highest expression and main function in unipotent Ly6C^+^ CD115^−^ granulocyte progenitors (GPs) and Ly6C^+^ CD115^+^ MPs downstream of GMPs^49^, we next evaluated the composition of the GMP population in the fetus and the adult with regards to these two markers. In contrast to adult GMPs, where 60-70% of the GMP population expressed Ly6C, only 30% of the fetal GMP pool did (Figure 5D). Further subdivision of immunophenotypic GMPs into immature GMPs (Ly6C^−^ CD115^−^), GPs and MPs confirmed a significantly lower frequency of GPs and MPs and a significantly higher frequency of bipotent GMPs in E14.5 FL compared to ABM (Figure 5E), indicating that the ABM environment more efficiently drives terminal myeloid differentiation than does the FL. However, we did find that the FL GMP compartment contains a population of CD115^+^ cells that is comparable in frequency to that in ABM (Figure 5F) and that only half of these cells additionally express Ly6C, whereas the adult CD115^+^ GMP population almost exclusively consists of Ly6C^+^ cells (Figure 5G). To further characterize the fetal and adult CD115^+^ GMPs, we investigated the myeloid differentiation capacity of these cells. Strikingly, fetal CD115^+^ GMPs failed to give rise to any live progeny (Figure 5H), indicating that CD115^+^ GMPs in the FL represent terminally differentiated cells, despite half of the cells sharing the adult MP immunophenotype (Figure 5G). In stark contrast, 70-90% of the progeny of adult CD115^+^ GMPs expressed CD11b already after 5 days (Figure 5H), a far higher frequency than what we observed for unfractionated adult GMPs (Supplemental Figure 3G). The progeny of adult CD115^+^ GMPs exclusively consisted of monocytes, confirming that these cells represent unipotent MPs (Supplemental Figure 3I).

Taken together, our data show that the lower Irf8 expression in fetal relative to adult GMPs correlates with an impaired ability to generate fully mature Ly6C^+^ monocytes from fetal cells, as well as a diminished prevalence of Ly6C^+^ unipotent myeloid progenitors in the fetal GMP pool compared to the adult (Figure 5I). Our results further suggest that CD115^+^ GMPs in the fetus do not correspond to MPs in the adult, and that *in situ* GMP-derived monopoiesis is not yet fully developed in the E14.5 FL.

## Discussion

In this work, we have comprehensively described the cellular proteome of three subsets of lineage-biased HPCs in the fetus and adult, and characterized the cells’ function. Our functional assays should be regarded as a comparison of the cells’ capacities driven by intrinsic factors. For example, the monocyte output we observed from the adult CLPs do not fully represent their *in vivo* lineage potential, as previously reported^25^. Instead, by keeping the extrinsic factors of the *in vitro* culture experiments identical for fetal and adult cells we have been able to fully compare the cells’ differential functional potential caused by their intrinsic molecular differences.

Previous studies have reported more extensive myeloid potential and transcript-level multilineage priming in embryonic and fetal LMPPs compared to adult cells^18,23,52-54^. In contrast, our *in vitro* differentiation results and proteome data suggest a stronger association of adult LMPPs to the myeloid lineage. Direct comparisons to previous work are obstructed by the use of a multitude of different immunophenotypic definitions and lineage-potential assays across different studies, as well as discrepancies between transcript- and protein-level expression profiles. Our study underscores the necessity to perform proteomic analyses to fully understand the biology of HPCs and potential leukemia-initating cells. Indeed, the timing of the establishment of myelopoiesis has so far been left unresolved, possibly due to a lack of gene expression profiling on the protein-level^28,55^. Our data support the notion that HSC-derived myelopoiesis is not yet fully developed in the E14.5 FL and is subject to considerable ontogenic regulation^28^, which we have linked to differential expression of myelopoiesis-associated proteins. This includes that fetal GMPs express lower levels of Irf8 compared to their adult counterparts, and produce monocytes with a phenotype reminiscent of that observed in adult *Irf8−/−* mice^49,50^. Although we observed a significantly higher frequency of CD11b^−^ Ly6C^+^ CD115^+^ “MP-like” cells produced from fetal compared to adult GMPs *in vitro*, the FL did not contain an expanded MP population *in vivo*, in contrast to what has previously been reported for BM of mice deficient in Irf8^49^. A potential explanation for this is that the FL environment appears to less strongly promote terminal myeloid differentiation relative to the ABM environment. Investigations of the role of extrinsic factors are required to resolve this question.

Unexpectedly, while myosins are mostly associated with housekeeping roles, our data point towards an involvement of these proteins in regulating LMPP-derived myelopoiesis in the adult, and that granulopoiesis from adult LMPPs is partly dependent on myosin activity. These results support the previously reported leukemia-inhibitory effect of ROCK inhibition and MLC knockdown^42^. Further investigations are needed to resolve whether these effects are limited to adult blood cancers and would have little effect on *in utero*-derived leukemias.

Similarly to molecular differences between fetal and adult CMPs^28^, the expression of several proteins linked to a MegE fate were elevated in fetal GMPs relative to adult cells. However, this protein signature does not appear to confer the fetal cells with any substantial MegE potential, indicating that the proteins that are typically associated with MegE fate rather represent a more general inflammatory signature and that fetal GMPs share more features with effector cells than with multipotent progenitors. This is supported by our functional assays, which show that the capacity for mature myeloid cell generation is lower from fetal compared to adult GMPs.

Collectively, our work has uncovered several novel molecular features that distinguish fetal and adult lineage-biased HPCs and raised important questions regarding the developmental timing of mature blood cell production from different hematopoietic progenitor populations. Our findings provide a substantial contribution to the current understanding of fetal and adult hematopoiesis. We expect these results to aid in directing future research focused on the development of age-tailored therapeutic approaches for improving the clinical outcomes of infant and adult leukemia patients.

## Acknowledgements

We greatly acknowledge the personnel at the FACS Core Facility at Lund Stem Cell Center as well as Sven Kjellström and Hong Yan at the BioMS Lund node for their expert technical assistance. We thank Parashar Dhapola for assistance with generating radar plots, and Fatemeh Safi and Charlotta Böiers for valuable input on experimental design. This work was supported by grants from the Swedish Research Council (to J.H., grant reference 2015-03063) and The Swedish Childhood Cancer Foundation (to J.H., grant reference TJ2106-0038).

## Authorship

J.H. and M.J. designed the study with input from E.S. and K.P.. M.J. performed experiments with assistance from K.P. and E.J.. M.J. analyzed data. J.H. supervised the study. M.J. prepared the figures and wrote the paper together with J.H. and input from all authors.

