## Supplemental methods and figures for "Ontogenic Shifts in Cellular Fate Are Linked to Proteotype Changes in Mouse Hematopoietic Progenitor Cells"

##### Mice

Wild-type C57Bl/6N mice purchased from Taconic Biosciences or bred in-house were used for all experiments. All experiments involving animals were performed in accordance with ethical permits approved by the Swedish Board of Agriculture. Animals were housed in individually ventilated cages (IVC) and provided with sterile food and water *ad libitum*. Adult mice used in experiments were 6-10 weeks old. E14.5 embryos were obtained by timed pregnancies overnight. The morning after mating was considered E0.5. For proteome analysis, equal numbers of male and female mice were used.

##### Flow cytometry and FACS

Adult BM was extracted from hind limbs, hip bones, forelimbs, shoulders, sternum and spine collected in Hank's Balanced Salt Solution (HBSS; HyClone). Single-cell suspension of adult BM was obtained by crushing bones using a mortar and pestle and passing cell suspensions through a 40 μm filter. Red blood cells were lysed by briefly incubating adult BM cells with ammonium chloride solution (StemCell Technologies) on ice. Lineage-positive cells were removed from adult BM cell suspension by depletion for Gr-1 (RB6-8C5; BioLegend), Ter119 (TER119; BioLegend), CD3 (145-2C11; BioLegend), B220 (RA3-652; BioLegend) and CD11b (M1/70; BioLegend) using biotin-conjugated antibodies with MACS anti-biotin beads using the autoMACS Pro Separator (Miltenyi Biotech) or LS columns (Miltenyi Biotech). FLs were extracted from embryos collected at E14.5 gestation. Single-cell suspension of FL was obtained by mechanical dissociation and passing through a 40 μm filter. Red cells were removed from FL cell suspensions by depletion for Ter119 using a biotin-conjugated antibody with MACS anti-biotin beads using the autoMACS Pro Separator or LS columns. FL and adult BM were surface-stained with fluorophore-conjugated antibodies against either Sca-1 (D7; BioLegend), c-Kit (2B8; BioLegend), Gr-1, Ter119, CD3, B220, Flt3 (A2F10; BioLegend), IL-7Rα (A7R34; BioLegend) and CD150 (TC15-12F12.2; BioLegend) (LMPP/CLP cocktail), or Sca-1, c-Kit, Gr-1, Ter119, CD3, B220, CD41 (MWReg30; BioLegend), CD150 and CD16/32 (93; BioLegend) (GMP cocktail). Anti-Ly6c (HK1.4; BioLegend), -CD115 (AFS98; BioLegend) and -CD63 (NVG-2; BioLegend) were included in the GMP antibody cocktail for experiments where the GMP population was resolved into MPs and GPs. Adult cells were additionally stained for CD11b. In all flow cytometry and FACS experiments, cells were incubated with 7AAD (Merck) briefly before analysis to exclude dead cells. All flow cytometry and FACS experiments were performed on BD FACSAriaIIu (70 μm nozzle), BD FACSAriaIII (70 μm nozzle), BD LSRFortessa or BD LSRFortessa X-20 instruments at the FACS Core Facility at Lund Stem Cell Center. Data analysis was performed in FlowJo (BD).

##### Sample preparation for proteome analysis

FACS-sorted cells were collected in ice-cold HBSS, centrifuged and stored as dry pellets at -80 °C until further use. Pellets corresponding to 100,000 cells were processed using in-StageTip (iST) NHS sample preparation kit (PreOmics) in accordance with manufacturer's protocol. Digested peptides were labeled using TMT6plex reagents (Thermo Scientific) in accordance with manufacturer's protocol. Immediately before use, TMT6plex reagents were equilibrated to room temperature. Vials containing 0.8 mg of TMT label were dissolved in 41 μl of anhydrous acetonitrile (ACN). Labeling was performed by addition of 10 μl dissolved TMT6plex to each sample and incubation for 1 hour at RT. Labeling reagents were swapped in the second and third biological replicate. Following desalting, labeled peptides were combined and dried by vacuum centrifugation. High-pH-reverse phase (HpH-RP) pre-fractionation was carried out as previously described[1] with some modifications. HpH-RP columns were assembled by packing C18-AQ beads onto 4 layers of C8 membrane in a 200 μl pipette tip via centrifugation. Columns were conditioned and equilibrated by sequential addition of 100% methanol followed by 80 and 20.8% ACN in ammonium formate, and finally 100% ammonium formate. Dried peptides were dissolved in 100% ammonium formate and bound to the column by centrifugation. HpH-RP pre-fractionation was carried out by sequential elution into 13 fractions containing increasing concentrations of ACN in ammonium formate: 7.5% (F1), 9% (F2), 11% (F3), 12.5% (F4), 14.5% (F5), 17.5% (F6), 20.8% (F7), 25% (F8), 28% (F9), 30% (F10), 35% (F11), 40% (F12), and 80% (F13). To achieve an even distribution of peptides in LC-MS analysis, fractions were combined as follows: F1+F9, F2+F10, F3+F11, F4+F12, and F5+F13 (yielding a total of 8 fractions together with F6, F7 and F8 that were kept separate). Fractionated samples were dried by vacuum centrifugation and stored at -20 °C until further use. Prior to LC-MS analysis, samples were dissolved in 4% ACN/0.1% formic acid (FA).

##### LC-MS analysis

MS analyses were carried out on an Orbitrap Fusion Tribrid MS instrument (Thermo Scientific) equipped with a Proxeon Easy-nLC 1000 (Thermo Fisher) using a 120 min linear gradient separation followed by a synchronous precursor selection MS3 (SPS-MS3) method. Each fraction was injected twice. Injected peptides were trapped on an Acclaim PepMap C18 column (3 $\mu$m particle size, 75 $\mu$m inner diameter x 20 mm length, nanoViper fitting), followed by gradient elution of peptides on an Acclaim PepMap RSLC C18 100 Å column (2 $\mu$m particle size, 75 $\mu$m inner diameter x 250 mm length, nanoViper fitting) using 0.1% (v/v) FA in LC-MS grade water (solvent A) and 0.1% (v/v) FA in ACN (solvent B) as the mobile phases. Peptides were loaded with a constant flow of solvent A at 9 µl/min onto the trapping column and eluted via the analytical column at a constant flow of 300 nl/min. During the elution step, the percentage of solvent B was increased in a linear fashion from 5% to 10% in 2 minutes, followed by an increase to 25% in 85 minutes and finally to 60% in an additional 20 minutes. The peptides were introduced into the mass spectrometer via a Stainless Steel Nano-bore emitter (150 $\mu$m OD x 30 $\mu$m ID; 40 mm length; Thermo Scientific) using a spray voltage of 2.0 kV. The capillary temperature was set at 275 °C. Data acquisition was carried out using a data-dependent SPS-MS3 method. The full MS scan was performed in the Orbitrap in the range of m/z 380 to 1580 and at a resolution of 120,000 at full-width-half-max (FWHM) using an automatic gain control (AGC) of 4.0e5 and a maximum ion accumulation time of 50 ms. The top ten most intense ions selected in the first MS scan were isolated for ion trap collision-induced dissociation MS2 (CID-MS2) at a precursor isolation window width of 0.7 m/z, an AGC of 1.5e4, a maximum ion accumulation time of 50 ms and resolution of 30,000 FWHM. The normal collision energy was set to 35%. Precursor selection range for MS3 was set to m/z range 400–1200 in MS2. Orbitrap HCD-MS3 scans were acquired in parallel mode with synchronous precursor selection (ten precursors), normalized collision energy of 55% and a resolution of 15,000 FWHM in a range of 100–500 m/z., the fragment ion isolation width was set to 2 m/z, the AGC was 1.0e5 and the maximum injection time 120 ms.

#### MS data analysis and bioinformatic analysis

MS raw data were processed with Proteome Discoverer (version 2.2.0; Thermo Scientific). Enzyme was set to trypsin and a maximum of two missed cleavages were allowed. Cysteine acetylhypusinylation and N-terminal and lysine TMT6plex were set as static modifications. Methionine oxidation and N-terminal acetylation were set as dynamic modifications. The peak list was searched using the Sequest HT node against the Swissprot mouse data base (downloaded 2017.07.05; 25,320 protein entries) together with commonly observed contaminants and reversed sequences for all entries. An FDR of 1% was required for identification at both the peptide and the protein level. Precursor and fragment mass tolerance were set to 10 ppm and 0.6 Da, respectively. Unique and razor peptides were used for quantification. The co-isolation threshold was set to 75. Proteins were quantified based on the average corrected TMT reporter ion intensities from two technical replicates per sample. Data was normalized by adjusting the median of each channel to the median of the medians of the entire replicate. Principal component analysis (PCA) was performed on the average protein abundances for each cell type using Perseus (version 1.6.6.0). Correlation was assessed using the Pearson correlation coefficient. Statistical analysis was performed on ratios of normalized intensities using the Limma package in R/Bioconductor[2]. P values were corrected for multiple testing using Benjamini-Hochberg's method. Proteins with an adjusted p value < 0.05 were considered to be differentially expressed. The PANTHER classification system was used to retrieve classification of proteins[3]. For differentially expressed proteins, gene ontology (GO) enrichment analysis was performed using the functional annotation tool DAVID[4]. Redundant GO terms were filtered out using REVIGO[5]. Analysis of transcription factor (TF) expression was performed on proteins classified as TFs by PANTHER and validated as TFs by TRRUST[6]. Mapping of differentially expressed proteins to transcriptome data from BloodSpot[7] was carried out using the "normal mouse hematopoiesis" dataset. A radar plot was generated using min-max scaled median values of marker genes in each cell type.

##### OP9/OP9Dl1 co-culture assays

For coculture experiments, 50 fetal and adult HPCs were FACS-sorted into 48-well plates onto pre-established layers of 10,000 OP9/OP9DL1 cells per well. Cells were cultured in Opti-MEM + GlutaMax medium (Gibco) supplemented with 10% fetal calf serum (FCS; HyClone), 1% penicillin/streptomycin (Gibco) and 0.02% 50 mM 2-mercaptoethanol (Gibco), and 25 ng/mL stem cell factor (SCF), 25 ng/mL Flt3-ligand (Flt3l), 20 ng/mL interleukin (IL) -7, 10 ng/mL IL-6, 10 ng/mL granulocyte (G) -colony stimulating factor (CSF), 10 ng/mL granulocyte-macrophage (GM)-CSF and 10 ng/mL IL-3 (OP9), or 25 ng/mL SCF and 25 ng/mL Flt3l (OP9DL1). SCF was omitted after the first week of culture for OP9DL1 cocultures. All cytokines were purchased from Stem Cell Technologies. Prior to flow cytometric analysis, cultured cells were incubated with FC-block and stained with fluorophore-conjugated antibodies against NK1.1 (PK136; BioLegend), CD11c (N418; BioLegend), Ly6G (1A8; BioLegend), Ly6C, CD11b, CD19 (6D5 or 1D3/CD19; BioLegend), B220 and CD115, or CD25 (PC61; BioLegend), Thy1 (53-2.1; BioLegend), CD3, CD4 (RM4-5; BioLegend) and

CD8 (53-6.7; BioLegend). Anti-F4/80 (BM8; BioLegend) anti-Gr-1 (in place for Ly6G) were used when investigating the lineage-potential of fetal $CD41^-$ and $CD41^+$ GMPs.

#### Erythroid differentiation assay

Erythroid potential was evaluated by culturing 10 fetal and adult cells per well in Terasaki plates in X-VIVO 15 medium (Lonza) supplemented with 0.5% bovine serum albumin (BSA), 10% FCS, 100 U/mL penicillin/streptomycin, 2 mM L-glutamine (Gibco), 0.1 mM 2-mercaptoethanol, 50 ng/mL SCF, 50 ng/mL Flt3L, 20 ng/mL IL-3, 30 ng/mL thrombopoietin (TPO) and 5 U/mL erythropoietin (EPO, which was generously provided by Dr. Göran Karlsson). Erythroid colony formation was assessed by diaminofluorene (DAF) staining. .

#### Megakaryocyte differentiation assay

Fetal and adult GMPs were suspension cultured at 5000 cells per well in complete OptiMEM medium supplemented with 50 ng/mL TPO, 20 ng/mL IL-6 and 10 ng/mL IL-3. IFN$\alpha$ (3,125 U; Miltenyi Biotech) or sterile water was added to stress and control wells, respectively. Cells were collected after 16 hours and megakaryocyte potential was assessed using the MegaCult culture system (StemCell Technologies) in accordance with manufacturer's protocol.

#### Suspension culture of GMPs

Suspension cultures of fetal and adult myeloid progenitors were performed by culturing 5000 fetal and adult cells per well in complete OptiMEM medium supplemented with 25 ng/mL SCF, 10 ng/mL IL-6, 10 ng/mL IL-3, and 10 ng/mL macrophage (M)-CSF. Cells were cultured for 7-9 days. Cultured cells were incubated with FC-block and stained with fluorophore-conjugated antibodies against Ter119, CD3, B220, NK1.1, CD11c, CD11b, F4/80, CD115, Ly6G and Ly6C prior to flow cytometric analysis.

#### ROCK inhibition assays

50 fetal and adult LMPPs were co-cultured with OP9 cells as described above. ROCK inhibitor H1152 (2 $\mu$M; R&D systems) or sterile water was added to wells at the start of culture and again 24 hours later. Media was exchanged after 48 hours to remove any remaining H1152. For experiments involving IFN$\alpha$, cells were cultured in the presence of IFN$\alpha$ (31.25 U) or sterile water for 18 hours prior to media exchange and addition of H1152. Cells were cultured for an additional 4-12 days before analysis. Cultured cells were incubated with FC-block and

stained with fluorophore-conjugated antibodies against NK1.1, Gr-1, CD11b, CD19, B220 and CD5 (53-7.3; BD) prior to flow cytometric analysis. Cell number and viability was assessed with trypan blue using an automated cell counter (BioRad).

#### Statistical analysis

Differences between two groups were assessed by two-tailed Students' t-test using Prism software version 8 (GraphPad software). Error bars represent SD. ****$p<0.0001$, ***$p<0.001$, **$p<0.01$, and *$p<0.05$.

#### Data sharing

Proteome data has been deposited to the MassIVE repository.

#### Supplemental tables and legends

**Supplemental Table 1. Proteome of fetal and adult LMPPs, CLPs and GMPs.** Data for all identified proteins with protein FDR confidence $q<0.01$ ('High')

### Supplemental figures and legends

#### Supplemental Figure 1

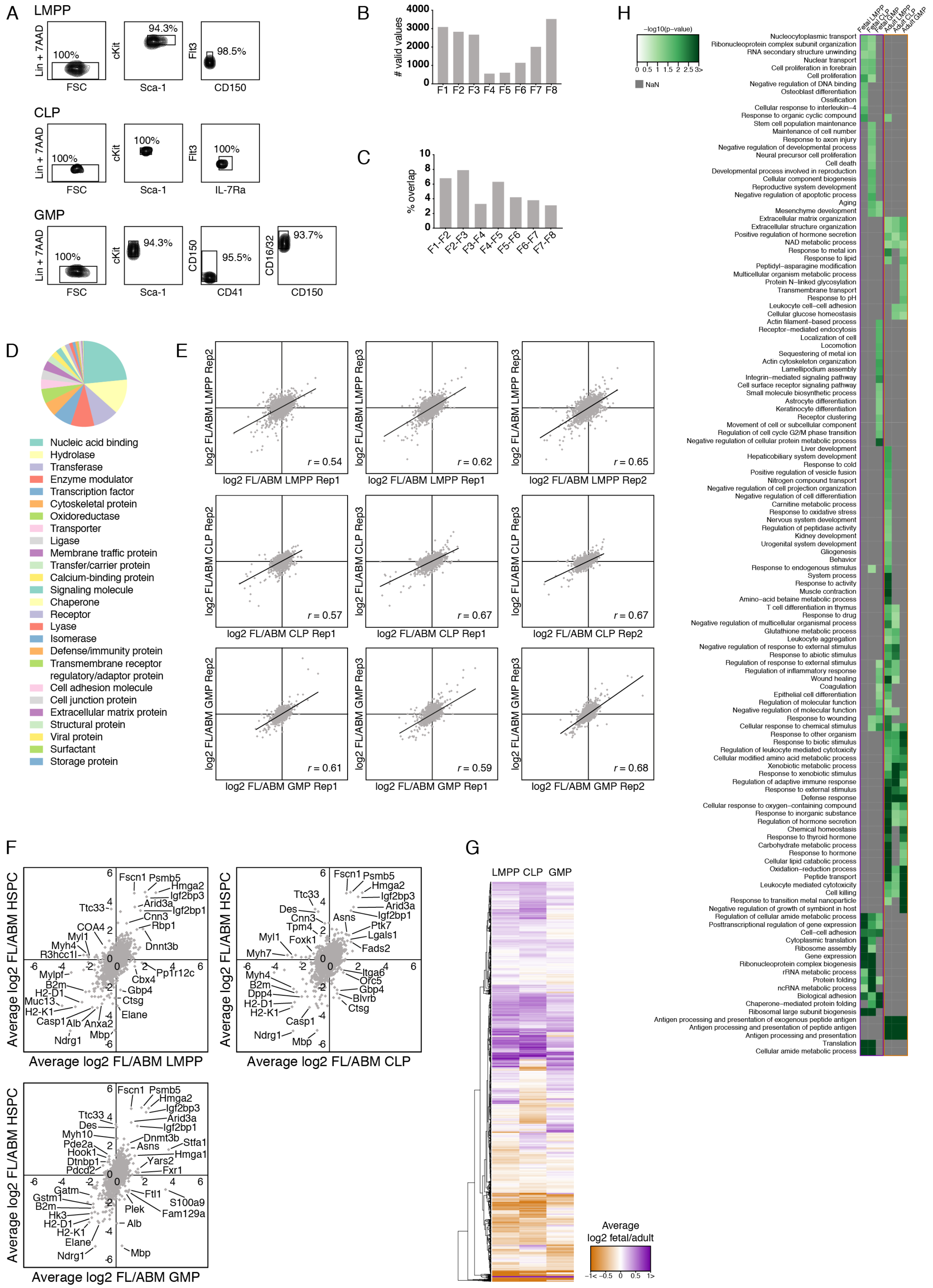

**Supplemental Figure 1. Comprehensive protein-level characterization of fetal and adult LMPPs, CLPs and GMPs.** (A) Representative FACS plots showing purity of sorted LMPPs, CLPs and GMPs. (B, C) Number of peptides quantified in each of the eight HpH-RP fractions analyzed by MS (B) and overlap between peptides identified in adjacent fractions (C) from six replicates of 100,000 FACS-sorted LS$^-$K cells. (D) PANTHER classification of the 4189 identified proteins. Proteins from 26 different classes were identified. (E) Protein expression correlation between the three replicates shown as the Pearson correlation coefficient (*r*). (F) Correlation between fetal/adult LMPP, CLP and GMP protein ratios and fetal/adult HSPC protein ratios[1]. (G) Heatmap showing average expression of 4021 proteins quantified in both the fetus and the adult in all three cell types. (H) Heatmap depicting enrichment (-log10 p-value) of all biological processes identified as enriched among differentially expressed proteins in any cell type. Only processes with a p-value < 0.05 are shown. Processes that were not detected are depicted in grey.

Supplemental Figure 2

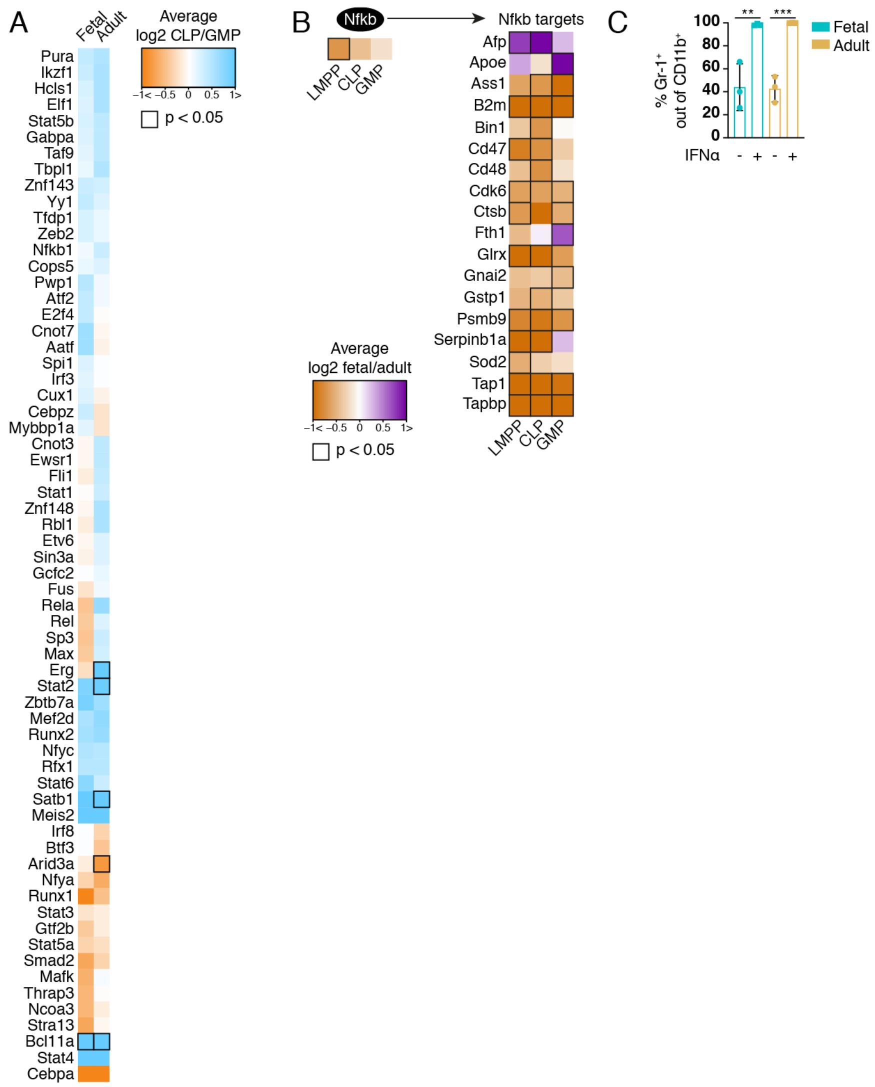

**Supplemental Figure 2. Protein expression differences indicate differential functionality of fetal and adult HPCs.** (A) Heatmap depicting the relative expression of proteins classified as transcription factors by PANTHER[52] and validated by TRRUST[53]. Proteins differentially expressed between CLPs and GMPs are indicated by a black frame. Data represent the average of three biological replicates. (B) Protein expression of Nfκb and its targets in fetal and adult LMPPs, CLPs, and GMPs. Differentially expressed proteins are indicated by a black frame. (C) Frequency of Gr-1$^{+}$ granulocytes derived from LMPPs treated with IFNα. Error bars are ± SD. $**p<0.01$, $***p<0.001$.

Supplemental Figure 3

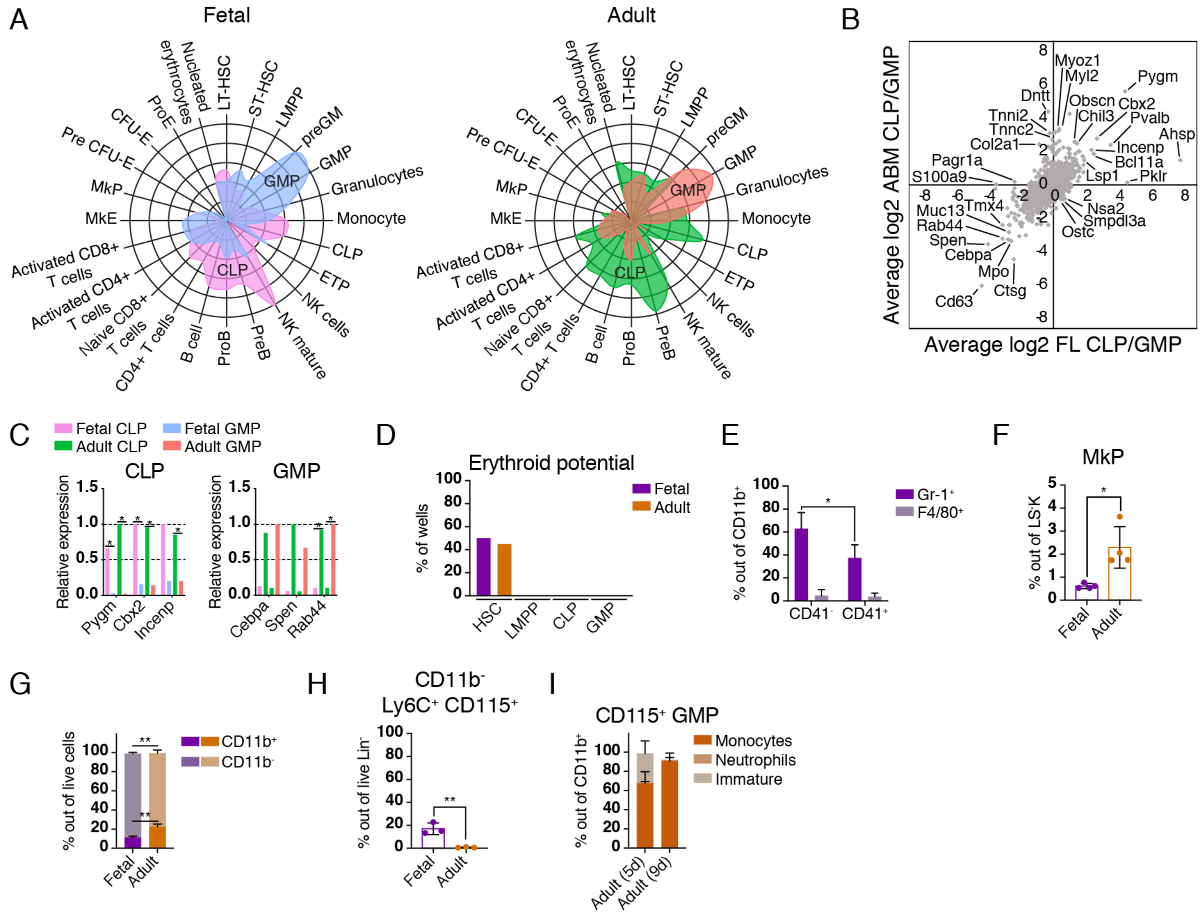


**Supplemental Figure 3. Expression of lineage-associated proteins predicts potential of fetal and adult HPCs.** (A) Radar plots depicting the association of differentially expressed proteins in CLPs and GMPs with known transcriptional profiles of murine hematopoietic cell subsets. (B) Correlation between fetal CLP/GMP and adult CLP/GMP protein ratios. (C) Relative expression of proteins identified as CLP and GMP signature proteins in (B). Data represent the average of three biological replicates. (D) Erythroid potential in fetal and adult LMPPs, CLPs and GMPs.

HSCs were included as a positive control. (E) *In vitro* granulocyte (NK1.1$^-$ CD11b$^+$ Gr-1$^+$ F4/80$^-$) and macrophage (NK1.1$^-$ CD11b$^+$ Gr-1$^-$ F4/80$^+$) potential of fetal CD41$^-$ and CD41$^+$ GMPs. (F) Frequency of MkPs (LS$^-$K CD41$^+$ CD150$^+$) in E14.5 FL and ABM. (G) Frequency of CD11b$^+$ and CD11b$^-$ cells derived from fetal and adult GMPs. Data are depicted as mean of three biological replicates. (H) Frequency of CD11b$^-$ Ly6C$^+$ CD115$^+$ cells immunophenotypically analogous to MPs derived from fetal and adult GMPs. (I) Frequency of neutrophils, monocytes and immature myeloid cells derived from adult CD115$^+$ GMPs. Data are depicted as mean of three biological replicates. Error bars are ± SD. *$p<0.05$, **$p<0.01$.
